# Automatic, web-based typing of *Clostridioides difficile* ribotypes using MALDI-TOF MS

**DOI:** 10.1101/2024.01.24.576988

**Authors:** Mario Blázquez-Sánchez, Alejandro Guerrero-López, Ana Candela, Albert Belenguer-Llorens, José Miguel Moreno, Carlos Sevilla-Salcedo, María Sánchez-Cueto, Manuel J. Arroyo, Mark Gutiérrez-Pareja, Vanessa Gómez-Verdejo, Pablo M. Olmos, Luis Mancera, Patricia Muñoz, Mercedes Marín, Luis Alcalá, David Rodríguez-Temporal, Belén Rodríguez-Sánchez, the AutoCdiff Study Group, Mariela Martínez, Cristina Verdú, Juan Soliveri, Juan Cuadros, Felipe Pérez, Alberto Nieto, Luís Moisés Ruiz, María Simón, Joan López, Óscar Cuevas, Carmen Ezpeleta, Soledad Illescas

## Abstract

*Clostridioides difficile* is a major cause of hospital-acquired diarrhea, posing significant clinical challenges due to its high morbi-mortality rates and its involvement in nosocomial outbreaks. Detecting its toxigenic ribotypes rapidly and accurately is crucial for effective outbreak control. This study aimed to create a rapid diagnostic methodology based on MALDI-TOF MS and Machine Learning algorithms to differentiate toxigenic *C. difficile* RTs.

MALDI-TOF spectra were acquired from 379 clinical isolates sourcing from 10 Spanish hospitals and analysed using Clover MSDAS, a specific software for MALDI-TOF spectra analysis, considered as the state-of-the-art tool for this purpose, and AutoCdiff, an *ad hoc* software developed in this study.

Seven biomarker peaks were found to differentiate epidemic RT027 and RT181 strains from other RTs (2463, 3353, 4933, 4993, 6187, 6651 and 6710 *m/z*). Two peaks (2463 and 4993 *m/z*) were specifically found in RT027 isolates while combinations of the other 5 peaks allowed the differentiation of RT181 from other ribotypes. Automatic classification tools developed in Clover MSDAS and AutoCdiff using the specific peaks and the entire protein spectra, respectively, showed up to 100% balanced accuracy. Both methods allowed correct ribotype assignment for isolates sourcing from real-time outbreaks.

The developed models, available from Clover MSDAS and the AutoCdiff website -https://bacteria.id-offer researchers a valuable tool for quick *C. difficile* ribotype determination based on MALDI-TOF spectra analysis. Although further validation of the models is still required, they represent rapid and cost-effective methods for standardized *C. difficile* ribotype assignment.

## INTRODUCTION

*Clostridioides difficile*, a Gram-positive, anaerobic, spore-forming, faecal-oral transmitted rod, is the leading cause of infectious, antibiotic-related, nosocomial and community-acquired diarrhoea (1, 2). Ribotype 027 (RT027) is of great clinical interest and has been reportedly associated with hospital outbreaks (3, 4). Strains belonging to RT027 have been considered as “hypervirulent” due to the hyperproduction of toxins (2). Other ribotypes (RT), such as the hypervirulent RT078 and RT181, have recently been reported as highly prevalent (9.0%) and emerging in European countries, respectively (4, 5).

The Xpert^®^ *C. difficile* BT assay has been implemented worldwide for the rapid diagnostic of toxigenic *C. difficile* strains from faeces (6). This test is fast -50 min from a direct sample- and simple to perform (7). It allows the detection of toxin B (*tcdB*), binary toxin (*cdtA*) and the deletion in position 117 of the regulatory gene *tcdC* associated to RT027, but present in other *C. difficile* RTs. Therefore, the targeted genes do not allow the differentiation of RTs closely related to RT027, such as RT181 or RT176, belonging to clade 2 ST1. The Xpert^®^ *C. difficile* BT assay classifies these RTs as “presumptive RT027” (8, 9).

Matrix-Assisted Laser Desorption/Ionization Time of Flight (MALDI-TOF) Mass Spectrometry (MS), coupled with Machine Learning (ML) algorithms for spectra analysis, represents an efficient alternative to DNA-based methods -Pulse Field Gel Electrophoresis and PCR ribotyping-for distinguishing *C. difficile* RTs rapidly -1h from bacterial culture- (10), with a cost per isolate below 1$ (11). However, the current state-of-the-art for this approach has notable limitations, lacking a standard procedure for spectra analysis and an accessible code or software.

The goal of this study was to develop an efficient methodology for specific typing of the hypervirulent *C. difficile* strains RT027 and RT181 utilizing MALDI-TOF MS and ML algorithms.

## METHODS

### Bacterial isolates

A total of 379 clinical isolates (Table 1) belonging to the most prevalent *C. difficile* RTs in Spain and Europe were included in this study (12). A subgroup of 348 isolates were obtained from faeces of patients with diarrheal symptoms sourcing from 10 Spanish hospitals. The samples were submitted to Hospital General Universitario Gregorio Marañón -HGUGM- for further characterization of the *C. difficile* RTs. The remaining 31 *C. difficile* isolates sourced from two suspected nosocomial outbreaks detected in Madrid during May-June 2023 in the Hospital Central de la Defensa Gómez Ulla -HCDGU- (n=28) and HGUGM (n=3). All isolates were analyzed by Xpert^®^ *C. difficile* BT assay -Cepheid, USA- (6). Subsequently, they were characterized by PCR-ribotyping (13-15). Briefly, the intergenic spacer region located between the 16S rRNA and 23S rRNA encoding regions was amplified by PCR. The DNA fragments were analysed by capillary electrophoresis. Phylogenetic analysis of the ribotyping profiles was performed using the Bionumerics 7.2 software (bioMérieux, Marcy l’Etoile, France) and WebRibo (https://webribo.ages.at/) (13).

**Table 1.**
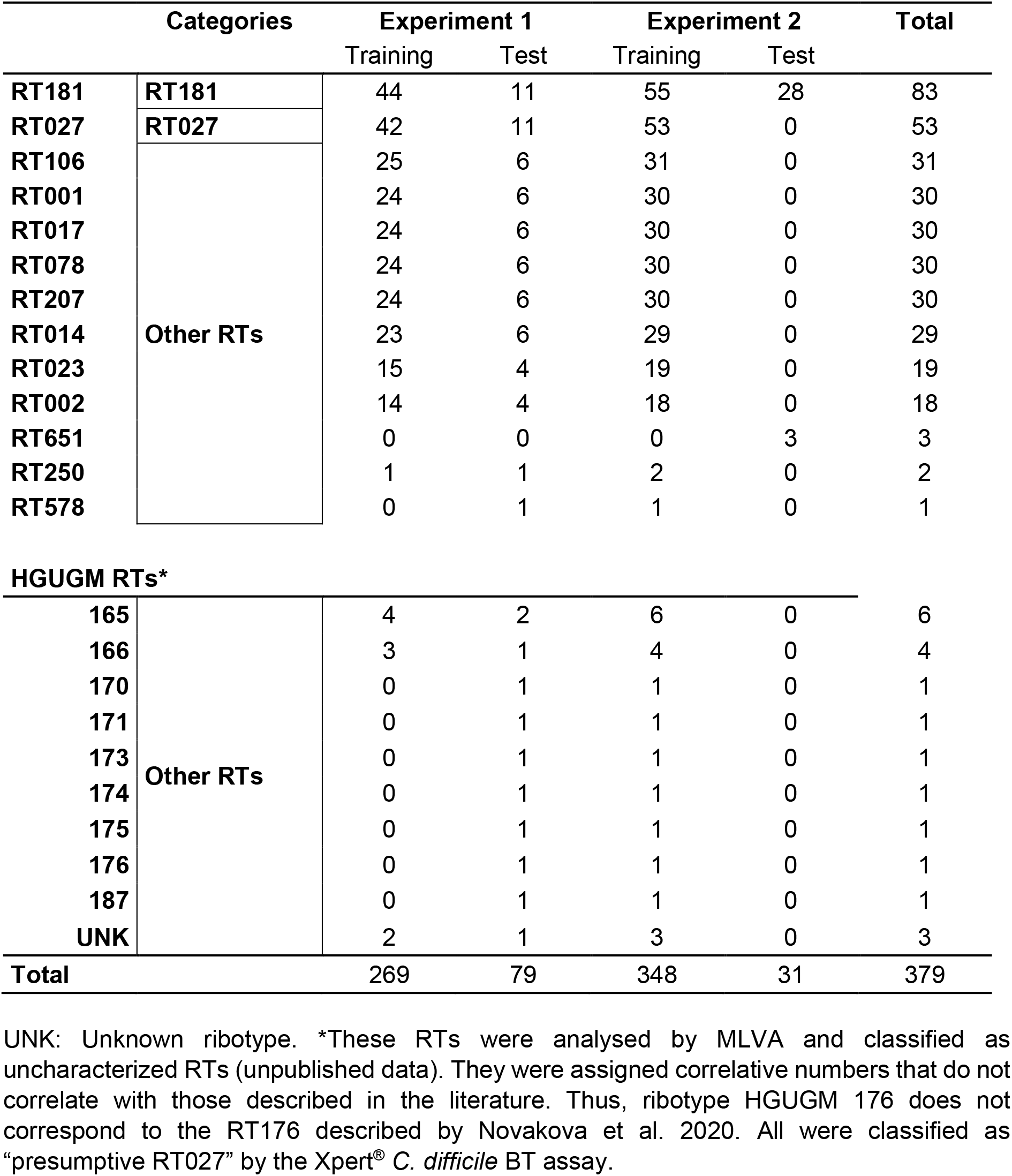
List of *C. difficile* isolates, classified according to their respective ribotype (RT) and the classification model in which they were included.

Prior to MALDI-TOF MS analysis, samples were plated on Brucella Blood Agar (Beckton Dickinson^®^, Franklin Lakes, NJ, US) and incubated under anaerobic conditions at 37ºC during 48h. Before analysis, all isolates were re-identified using MALDI-TOF MS in an MBT Smart MALDI Biotyper (Bruker Daltonics, Bremen, Germany).

### MALDI-TOF MS spectra acquisition

For the acquisition of MALDI-TOF MS spectra, a small number of colonies of each *C. difficile* isolate was transferred to the MALDI plate in duplicates, overlaid with 1μl of 100% formic acid, allowed to dry and covered with 1μl of organic HCCA (α-Cyano-4-hydroxycinnamic acid) matrix. Two spectra were acquired from each spot in the positive ion mode within the 2,000 to 20,000 Da range. Consequently, for each bacterial isolate, four spectra were available, consistent with the conditions applied for spectra analysis (16).

### Spectra preprocessing

Clover MSDAS applied the following pre-processing pipeline: 1) Square root variance stabilization; 2) Smoothing via Savitzky-Golay filter (window length: 11; polynomial order: 3); 3) Baseline subtraction using Top-Hat filter (factor 0.02); 4) Peak alignment with constant mass tolerance of 2 Da and linear mass tolerance of 600 ppm and 5) TIC normalization. On the other hand, AutoCdiff applied this pre-processing pipeline: 1) Square root variance stabilization; 2) Smoothing via Savitzky-Golay filter (window length: 11; polynomial order: 3); 3) Baseline subtraction using Top-Hat filter (factor 0.02); and 4) TIC normalization (17-19). Replicates were not merged but treated as unique spectra in both software packages, to mimic a real-time scenario.

### Automatic ML bacteria typing

Clover MSDAS, a commercial *ad hoc* software developed by Clover Bioanalytical Software (Granada, Spain, https://cloverbiosoft.com) was applied for biomarker detection and automatic typing with well-known ML algorithms: Random Forest (RF), K-Nearest Neighbor (KNN), Support Vector Machine (SVM) and Light Gradient Boosting Machine (Light-GBM). In Clover MSDAS, all algorithms underwent training using biomarker peaks identified by the software, as described in previous studies (20-22). The use of these biomarkers for differentiation of specific categories is referred to as Expert-Knowledge (EK). The extraction of EK involved conducting a univariate analysis in Clover MSDAS, utilizing peak intensities to assign binary labels for Area Under the Curve (AUC) calculations. Peaks yielding an AUC greater than 0.7 were subsequently selected as potential biomarkers.

On the other hand, AutoCdiff is a novel, open-source tool developed in this study, designed to streamline the process of automatic bacteria typing. It features pre-built Bayesian ML models, including algorithms like the Factor Analysis-Variational AutoEncoder (FA-VAE) (23) and Dual Bayesian Logistic Regression with Feature Selection (DBLR-FS) (24), as well as foundational models such as RF and Decision Trees (DT). In AutoCdiff, algorithms use the complete spectrum for input, unrestricted to specific biomarkers. An additional approach for DBLR-FS allows the incorporation of EK as input to speed up the model convergence. Consequently, biomarkers extracted by Clover MSDAS were utilized as Bayesian priors for the model’s weights, thereby enriching the model with EK. AutoCdiff has been allocated in a webpage (https://bacteria.id) to allow MALDI users to efficiently analyze MALDI-TOF MS spectra by simply uploading raw data.

Regarding cross-validation Clover MSDAS utilized fixed parameters: RF with a minimum of 2 isolates per split, 1 leaf, 100 estimators, and sqrt maximum features; SVM with a linear kernel; Light-GBM with 32 leaves, a 0.1 learning rate, and 100 estimators and KNN with 5 neighbors. In AutoCdiff, a 10-fold cross-validation was applied for DT and RF algorithms, experimenting with various parameters like maximum depth -2, 4, 6, 8-g, minimum isolates per split -2, 4, 6-, minimum leaves -1, 2, 4-, and maximum features -auto, sqrt, log2-. The FA-VAE and DBLR-FS models, due to their probabilistic nature, bypassed the need for cross-validation, enabling direct estimation of model hyperparameters and providing as output probabilities and uncertainties.

### Experiment design

To evaluate the accuracy of the available ML algorithms, two experiments were designed: i) to differentiate the main toxigenic RTs in the study collection, namely RT027 and RT181, directly from the MALDI-TOF MS spectra. To carry out this evaluation, a dataset (n=269) was built with *C. difficile* isolates from the most prevalent RTs in Spain and Europe (4, 12), such as RT001, RT014, RT106, RT207, RT023, RT002, and RT017 (Table 1). Isolates from this experiment were classified into three distinct categories: RT027 (n=44), RT181 (n=43) and “Other RTs” (n=182). To validate this model, an independent subset of strains (n=79) from the same 3 categories was used; ii) Finally, the accuracy of the ML algorithms was evaluated in a real-world scenario with isolates from two nosocomial outbreaks. All algorithms underwent training using the whole bacterial collection, encompassing 348 *C. difficile* isolates (Table 1). Subsequently, these algorithms were tested with isolates sourcing from two outbreaks detected in HCDGU (n=28) and HGUGM (n=3) in real time (Figure S1).

### Ethics Statement

This study was approved by the University Hospital Gregorio Marañon Ethics Committee (CEIm) -Study number MICRO.HGUGM.2021-025-. The Committee considered that all the conditions for waiving informed consent were met since the study was conducted with microbiological samples and not with human products.

## RESULTS

### Biomarker Peak Analysis

A total of 7 potential biomarker peaks were identified (Table 2) using Clover univariate analysis. Specific peaks for RT027 differentiation were found at 2463 and 4933 *m/z* (Figure 1). A peak at 4993 *m/z* was absent in RT027 but present in RT181 and other RTs while peaks at 3353, 6187, 6651 and 6710 *m/z* were shown to differentiate both RT027 and RT181 from Other RTs (Figure 1). Three of these peaks, at 3353 -the same peak as 6710 *m/z* with double charge-, 6651 and 6710 *m/z*, were correlated with two isoforms of the methionine-cleaved ribosomal protein L28, using the Ribopeaks database (https://lcad.deinfo.uepg.br/~ppgca/ribopeaks/).

**Table 2.** Biomarker peaks selected for the construction of classification models. Area Under the Curve values are represented in this table.

| *m/z* | 2463 | 3353 | 4933 | 4993 | 6187 | 6651 | 6710 |
| --- | --- | --- | --- | --- | --- | --- | --- |
| RT027 | 0.93 | 0.85 | 0.98 |  | 0.88 |  | 0.96 |
| RT181 |  | 0.90 |  | 0.98 | 0.89 |  | 0.98 |
| Other RTs |  |  |  | 0.77 |  | 0.97 |  |

RT: Ribotype.

**Figure 1.**
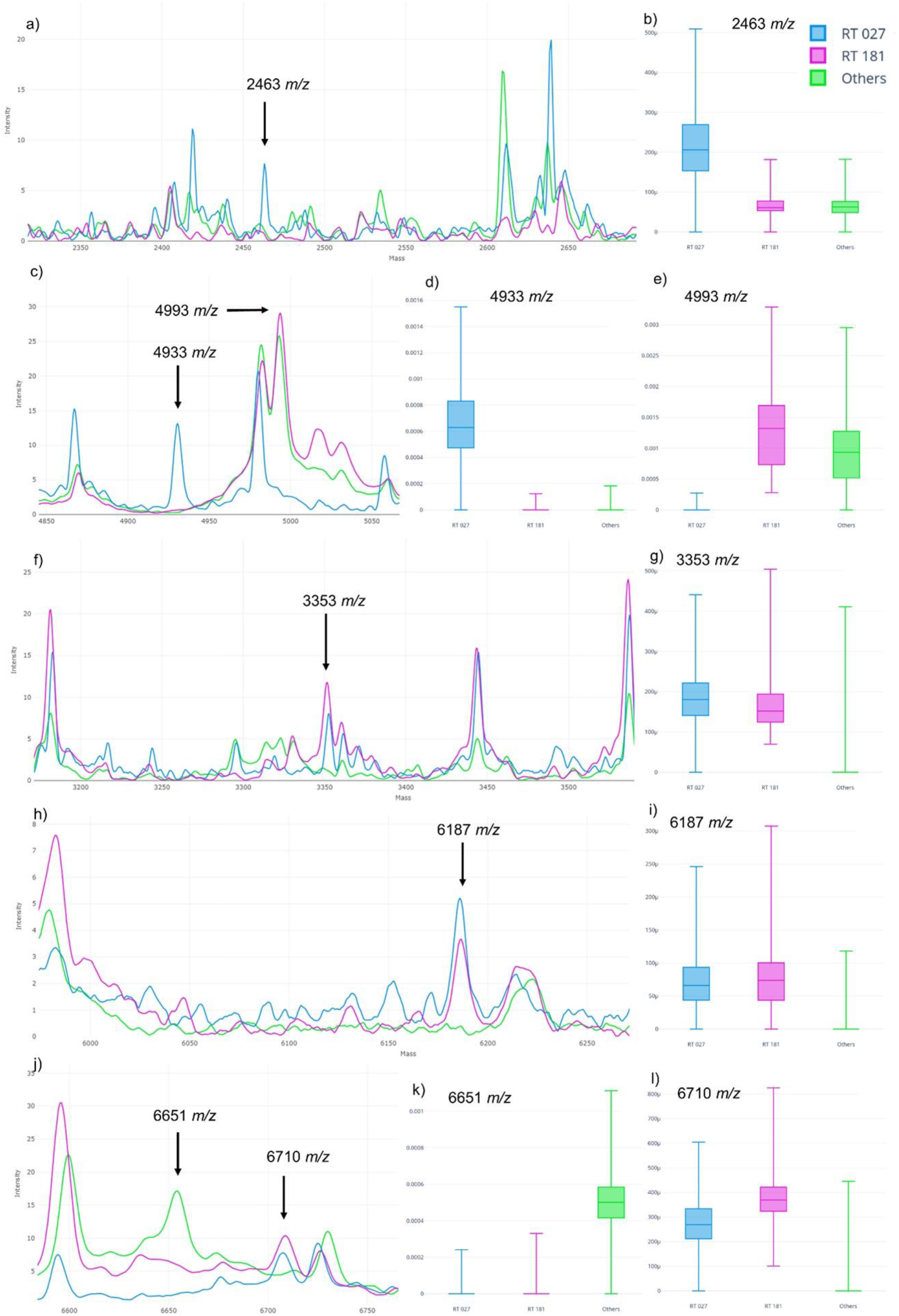
Peaks used for creation of predictive models for Clover Mass Spectrometry Data Analysis Software. a) Peak 2463 *m/z*. b) Intensities of 2463 *m/z* according to model categories. c) Peak 3353 *m/z*. d) Intensities of 3353 *m/z*. e) Peaks 4933 and 4993 *m/z*. f) Intensities of 4933 *m/z*. g) Intensities of 4993 *m/z*. h) Peak 6187 *m/z*. i) Intensities of 6187 *m/z*. j) Peaks 6651 and 6710 *m/z*. k) Intensities of 6651 *m/z*. l) Intensities of 6710 *m/z*.

This was also suggested by the information about protein L28 for *C. difficile* available in the Uniprot database (https://www.uniprot.org/). The difference between these two isoforms is the aminoacidic substitution G9D according to Uniprot database (Table 3). Overall, these biomarker peaks were used as input for all Clover MSDAS algorithms and as a prior EK in the DBLR-FS algorithm of AutoCdiff.

**Table 3.** Two isoforms of ribosomal protein L28 in *Clostridioides difficile*, according to the ribotypes (RTs) included in this study. L28-M indicates the mass of the protein excluding the N-terminal methionine.

| Ribotypes | Mass L28 (Da) | Mass L28-M (Da) | Aminoacidic sequence of Uniprot database |
| --- | --- | --- | --- |
| RT027 and RT181 | 6837 | 6710 | MAKVCSVCDKGKVSGNQVSHSNKHNRKRTWSA<br>NLRSVRAIIDGAPKRVKVCTRCLRSGKIERA |
| Other RTs | 6779 | 6651 | MAKVCSVCGKGKVSGNQVSHSNKHNRKRTWSA<br>NLRSVRAIIDGAPKRVKVCTRCLRSGKIERA |

### Differentiation of *C. difficile* ribotypes

#### Experiment 1: RT027 and RT181 Specific Typing

We tested the capacity of the described methodology to differentiate RT027 from RT181 (a clinically relevant ribotype identified as “presumptive RT027” by Xpert^®^ *C. difficile* BT) and other “presumptive RT027” RTs.

Clover MSDAS algorithms used as input the 7 biomarkers listed in Table 2. Peaks at 4933 and 4993 *m/z* were the most significant ones in terms of intensity (Figure S2a). After applying PCA, three distinct clusters were obtained, correlated to the three defined categories -RT027, RT181, “Other RTs”- (Figure S2b). Regarding ML algorithms, RF yielded a balanced accuracy of 99.7%. The PPV of this model was 98.5% for RT027, 96.2% for RT181, and 97.3% for “Other RTs”. The balanced accuracy of RF model was followed by KNN with 96.7% and SVM with 95.6% (Table 4).

**Table 4.**
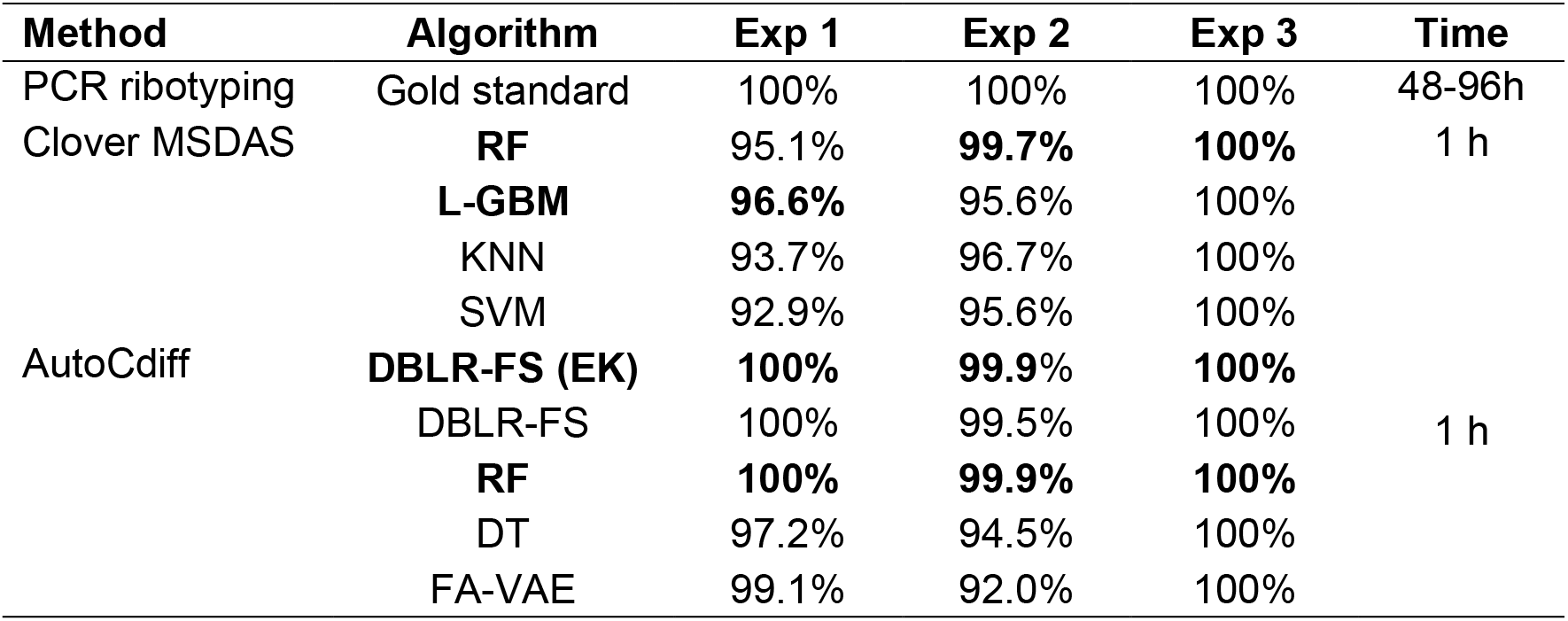
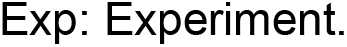
Balanced accuracy results on test sets for various experiments, comparing the performance of algorithms across Clover Mass Spectrometry Data Analysis Software (MSDAS) and AutoCdiff software packages. The Dual Bayesian Logistic Regression with Feature Selection (DBLR-FS) algorithm was evaluated with and without Expert Knowledge (EK) and all results are compared against the gold standard method for ribotype identification. In addition, time to obtaining results from positive culture is indicated.

| Method | Algorithm | Exp 1 | Exp 2 | Exp 3 | Time |
| --- | --- | --- | --- | --- | --- |
| PCR ribotyping | Gold standard | 100% | 100% | 100% | 48-96h |
| Clover MSDAS | **RF** | 95.1% | **99.7%** | **100%** | 1 h |
|  | **L-GBM** | **96.6%** | 95.6% | 100% |  |
|  | KNN | 93.7% | 96.7% | 100% |  |
|  | SVM | 92.9% | 95.6% | 100% |  |
| AutoCdiff | **DBLR-FS (EK)** | **100%** | **99.9%** | **100%** | 1 h |
|  | DBLR-FS | 100% | 99.5% | 100% |  |
|  | **RF** | **100%** | **99.9%** | **100%** |  |
|  | DT | 97.2% | 94.5% | 100% |  |
|  | FA-VAE | 99.1% | 92.0% | 100% |  |

Exp: Experiment.

In AutoCdiff, the algorithms used full spectra as input, incorporating EK-enhancement for DBLR-FS, as described in the methodology. Along the full spectra, DBLR-FS automatically detected 3 main peaks with high importance for RT differentiation: the presence of 4933 *m/z* for RT027 (Figure S3a), the absence of 4933 and 6651 *m/z* for RT181 (Figure S3b) and the presence of 6651 *m/z* combined with the absence of 6710 *m/z* for Other RTs (Figure S3c). These peaks correlated with those found by implementing the Clover MSDAS Biomarker Analysis. Algorithm DBLR-FS, with EK-enhancement, achieved a balanced accuracy of 99.9%, with PPV of 94.1% for RT027, 100% for RT181, and 100% for Others. Only one isolate, RT173 from the “Other RTs” group, was misclassified as RT027, likely due to the presence of the 4933 *m/z* peak, commonly associated with RT027.The inclusion of EK increased results from 99.5% to 99.9% while significantly speeding up training convergence.

Additionally, RF attained a balanced accuracy of 99.9%, while DT and FA-VAE exhibited slightly lower accuracies of 94.5% and 92.0%, respectively (Table 4).

#### Experiment 2: Application of the algorithms to real-time cases

The available algorithms were re-trained with the full dataset of spectra from our *C. difficile* collection (n=348) and tested during suspected nosocomial outbreaks at HCDGU (n=28) and HGUGM (n=3) in real-time. A proposal for implementation of this methodology in the clinical laboratory is showed in Figure 2.

**Figure 2.**
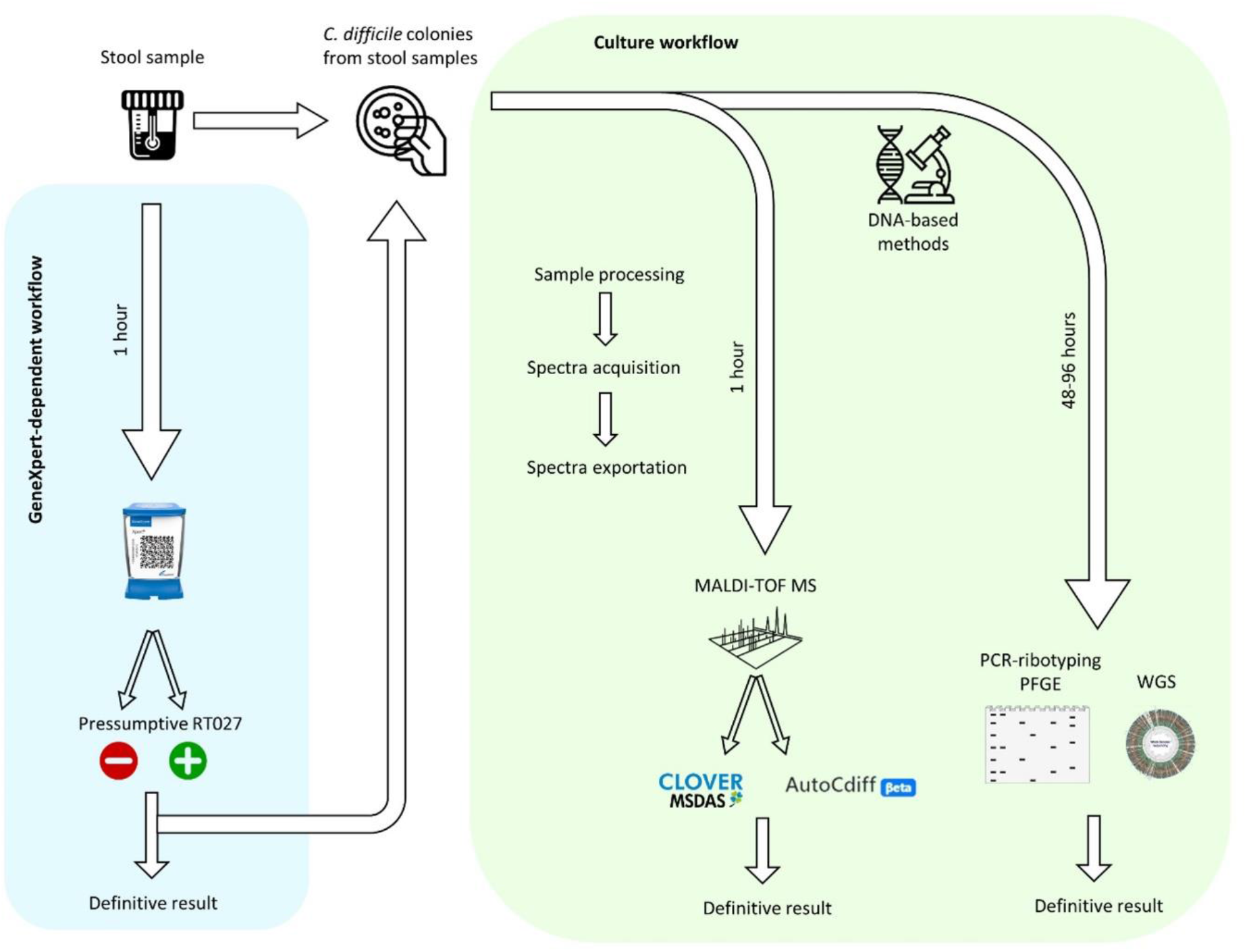
Proposed laboratory workflow with the implementation of *Clostridioides difficile* typing by MALDI-TOF MS methodology presented in this paper. The workflow shows a GeneXpert-dependent option and a direct culture option for those laboratories with no GeneXpert system available. PFGE: Pulsed-Field Gel Electrophoresis. WGS: Whole-Genome Sequencing.

All algorithms from both software packages consistently classified HCDGU isolates as RT181 and HGUGM isolates as “Other RTs” within 1 hour after culture positivity. PCR-ribotyping results validated the ML classifications 48h later, confirming that the outbreak *C. difficile* isolates from HCDGU belonged to RT181 and those from HGUGM corresponded to RT651 according to WebRibo database.

## DISCUSSION

In this study, it was demonstrated that the implementation of ML methods for the differentiation of *C. difficile* RTs based on MALDI-TOF MS spectra is a cost-effective, easy-to-apply, and reliable tool with great potential for rapid screening of these isolates, offering a real alternative to complex typing methods such as PCR ribotyping and PFGE. Its implementation could reduce the turn-around time for definitive characterization of the isolates to 48-72h hours from the reception of the clinical sample in the laboratory. The hands-on time of the method is approximately of 1 hour once the *C. difficile* isolates are grown on solid agar media.

A total of 7 biomarker peaks (2463, 3353, 4933, 4993, 6187, 6651, 6710 *m/z*) where found to differentiate RT027 from RT181 and Other RTs. Three of these biomarkers correlated with those automatically found by DBLR-FS (4993, 6651 and 6710 *m/z*) for the same purpose. Some of them have already been described in previous studies: protein peaks between 6647-6654 *m/z* and between 6707*-*6712 *m/z* (6651 *m/z* and 6710 *m/z* in this study, respectively) have been reportedly found in RT027 and RT176 (25, 26). These peaks could correspond to two isoforms of the ribosomal protein L28. In this study, the isoform 6710 *m/z* was found in RT027 and RT181, suggesting the great similarity between these two RTs, while the isoform 6651 *m/z* was specifically detected in the rest of the analyzed RTs, including those classified as “presumptive RT027”. Furthermore, the peak at 3353 *m/z* was found to be specific for RT027 and RT181, as well as the peak at 6710 *m/z*, suggesting the possibility that it is the same isoform of the L28 with double charge ions. However, no biomarker peak with double charge was found for the 6651 *m/z* isoform of the L28 protein.

The peak at 4928 *m/z* (4933 *m/z* in this study) has been described before as an uncharacterized protein present in RT027 (27). Our results correlated with this finding and showed that the 4933 *m/z* peak is specific to RT027 and the 2463 *m/z* peak, which we suggest may be the same protein with double charge. However, the peak at 4993 *m/z*, absent in RT027 and present in the other two remaining categories (“RT181” and “Other RTs”) had not been previously described, probably due to the limited literature about the RT181 available so far.

A recent paper applying Clover MSDAS reported a balanced accuracy >95.0% when differentiating hypervirulent *C. difficile* RTs (28). However, RT027 and RT176 could not be differentiated. Despite not having established a specific category for RT176, this study validates a methodology to differentiate RTs with characteristics like RT176 (“presumptive RT027” by Xpert^®^ *C. difficile* BT and cause of nosocomial outbreaks), as is the case of RT181. In this study, the algorithms Light-GBM and RF, available in Clover MSDAS, exhibited superior performance in distinguishing RT027 from other “presumptive RT027” isolates. This pattern was consistent across all algorithms, when differentiating RT027 and RT181 from the rest of the analyzed RTs. Furthermore, all algorithms achieved 100% accuracy in classifying the outbreak *C. difficile* isolates. In AutoCdiff, the DBLR-FS with EK and RF algorithms showed the highest accuracy in differentiating RT027 from other “presumptive RT027” isolates. The Bayesian methods in AutoCdiff, like DBLR-FS, do not need cross-validation and provide valuable uncertainty measurements, enhancing their practical utility. Therefore, the developed methodology has proven to be very useful to characterize C. difficile ribotypes in real time. It requires very limited expertise and means and favors the rapid implementation of control measures.

Although this study represents a step forward in automating MALDI-TOF MS data for bacterial typing, it still has limitations. The first one is the number of isolates analyzed, sourcing exclusively from Spain although from geographically distant hospitals. This limitation is expected to be overcome soon by the collaborations established with different research groups who have started to test the web-based classification models at www.bacteria.id (Figure S4). Besides, the need for a bacterial culture for MALDI-TOF MS analysis prolongs the detection period for 24-48h until colony growth, whereas the Xpert^®^ *C. difficile* BT can be performed from direct samples in a shorter time.

In conclusion, the integration of MALDI-TOF MS with ML methods presents a promising approach to effectively distinguish clinically significant *C. difficile* RTs, offering a time- and effort-effective screening solution, especially when an outbreak caused by hypervirulent *C. difficile* RTs is suspected. The availability of the developed methods could empower laboratories with limited molecular resources to enhance their diagnostic capabilities.

## FUNDING

This work is partially supported by grants PID2020-115363RB-I00, PID2021-123182OB-I00 and TED2021-132366B-I00 funded by MCIN/AEI/10.13039/501100011033, by the project PI18/00997 from the Health Research Fund (FIS. Instituto de Salud Carlos III. Plan Nacional de I+D+I 2013-2016) of the Carlos III Health Institute (ISCIII, Madrid, Spain) partially financed by the European Regional Development Fund (FEDER) ‘A way of making Europe’. The funders had no role in the study design, data collection, analysis, decision to publish, or preparation/content of the manuscript. AC (Rio Hortega CM21-00165), DRT (Sara Borrell CD22-00014) and BRS (Miguel Servet CPII19/00002) are funded by ISCIII. AGL and MBS are the recipients of the Intramural predoctoral contracts 2020 and 2022, respectively, from the Health Research Center of the Hospital Gregorio Marañón -IISGM-.

## Data Availability

Aligned with our dedication to the FAIR (Findable, Accessible, Interoperable, and Reusable) principles, we have made the dataset from this study publicly accessible on Zenodo. It can be found at https://doi.org/10.5281/zenodo.10370872.

## Code Availability

The code for all models implemented in AutoCdiff, which are open source, is freely accessible at our GitHub repository: https://github.com/aguerrerolopez/Clostridium. This repository includes comprehensive documentation and usage notes to facilitate replication and further exploration of our models.

## CONFLICTS OF INTEREST

The authors declare no conflict of interests. MJA and LM are employees of Clover Bioanalytical Software, S.L.

## Notes

https://doi.org/10.5281/zenodo.10370872.

### Competing Interest Statement

The authors have declared no competing interest.

### Summary of Updates

The results have been reorganized in a more logical manner: several classification models were trained for differentiation of the main C. difficile ribotypes involved in nosocomial outbreaks in our setting, i. e. RT027 and RT181, and validated with an external dataset. Then, the training and validation datasets were merged to train an improved model that was validated in real time with isolates from two hospitals. The models classified correctly 100% of the cases.

