## Supplemental Figures S1 to S4 for "Automatic, web-based typing of *Clostridioides difficile* ribotypes using MALDI-TOF MS"

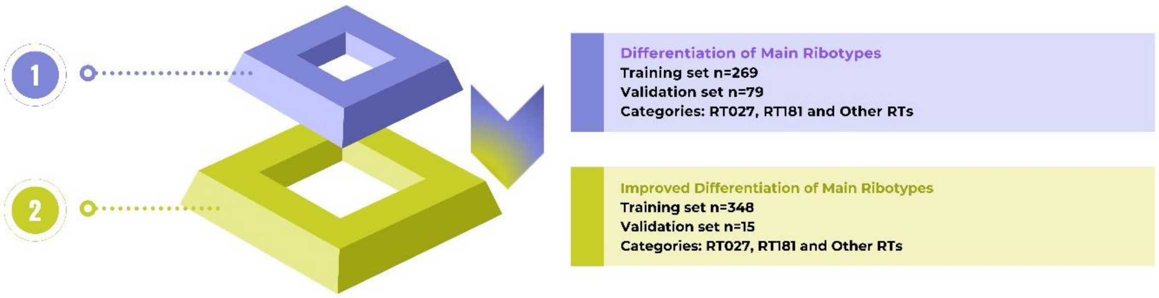


**Figure S1.** Description of the experiments developed for the classification of *C. difficile* protein spectra.

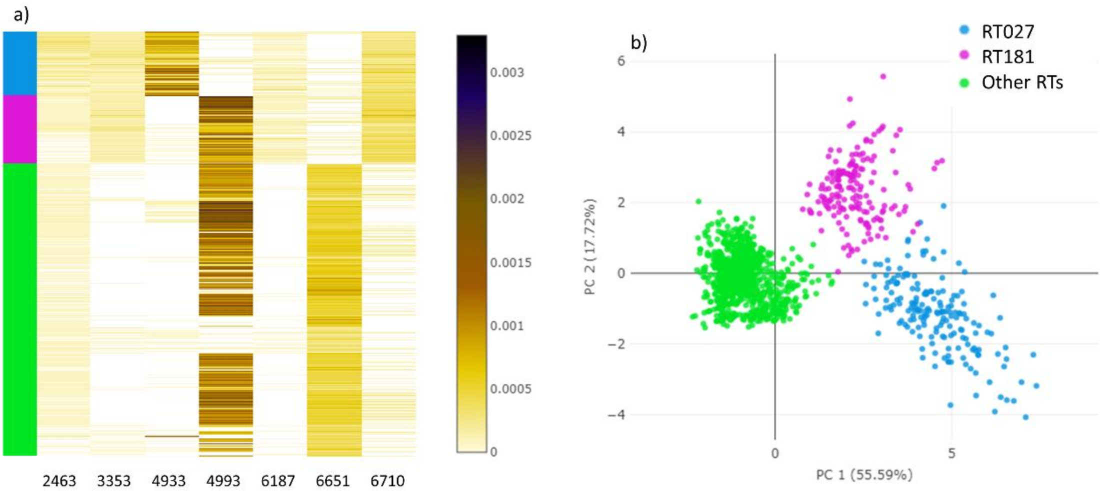


**Figure S2. a)** Intensity peak distribution heatmap of the seven selected peaks for Experiment 2. **b)** Principal Component Analysis using the peaks 2463, 3353, 4933, 4993, 6187, 6651 and 6710 *m/z* for differentiation of RT027 and RT181 isolates from isolates pertaining to Other ribotypes.

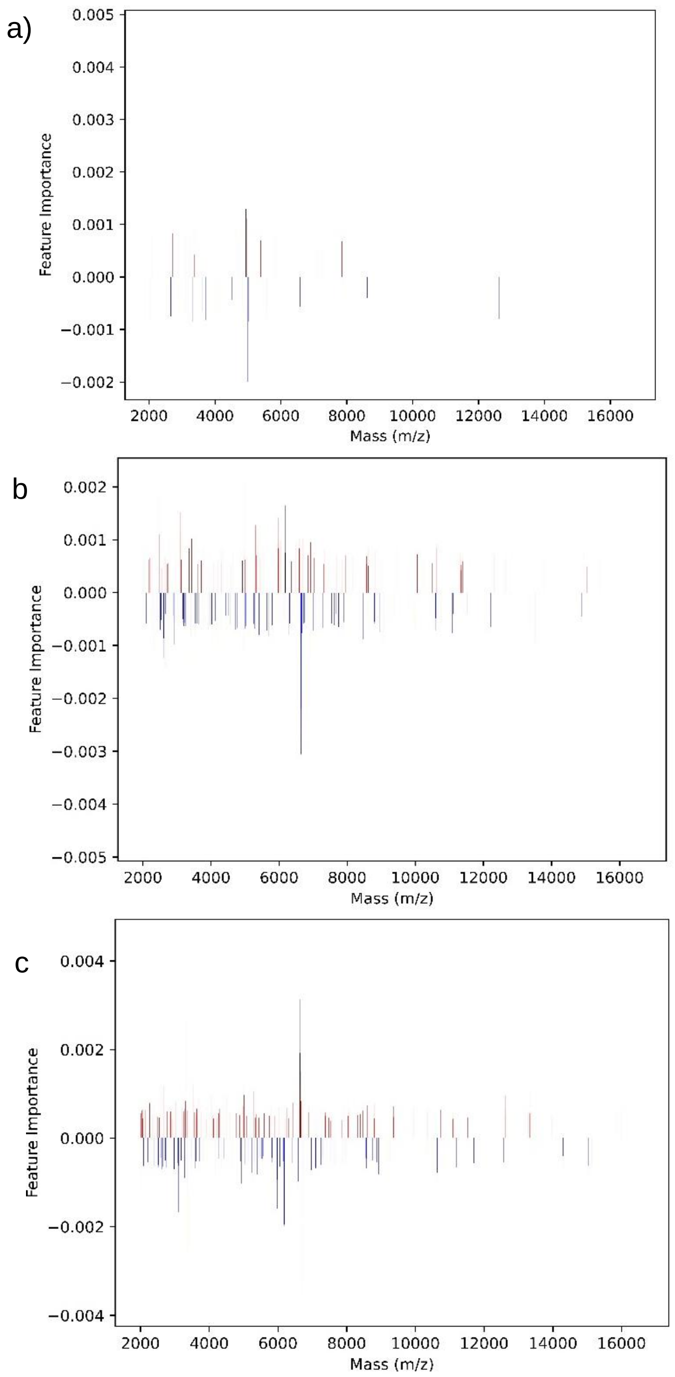


**Figure S3.** Feature importance of each protein mass obtained by DBL-FS algorithm. Positive weights are represented in red while negative ones in blue, additionally, the peaks introduced within the prior as Expert Knowledge (EK) are represented in black. **a)** Feature importances for ribotype (RT) 027. **b)** Feature importances for RT181. **c)** Feature importances for the category “Other RTs”.

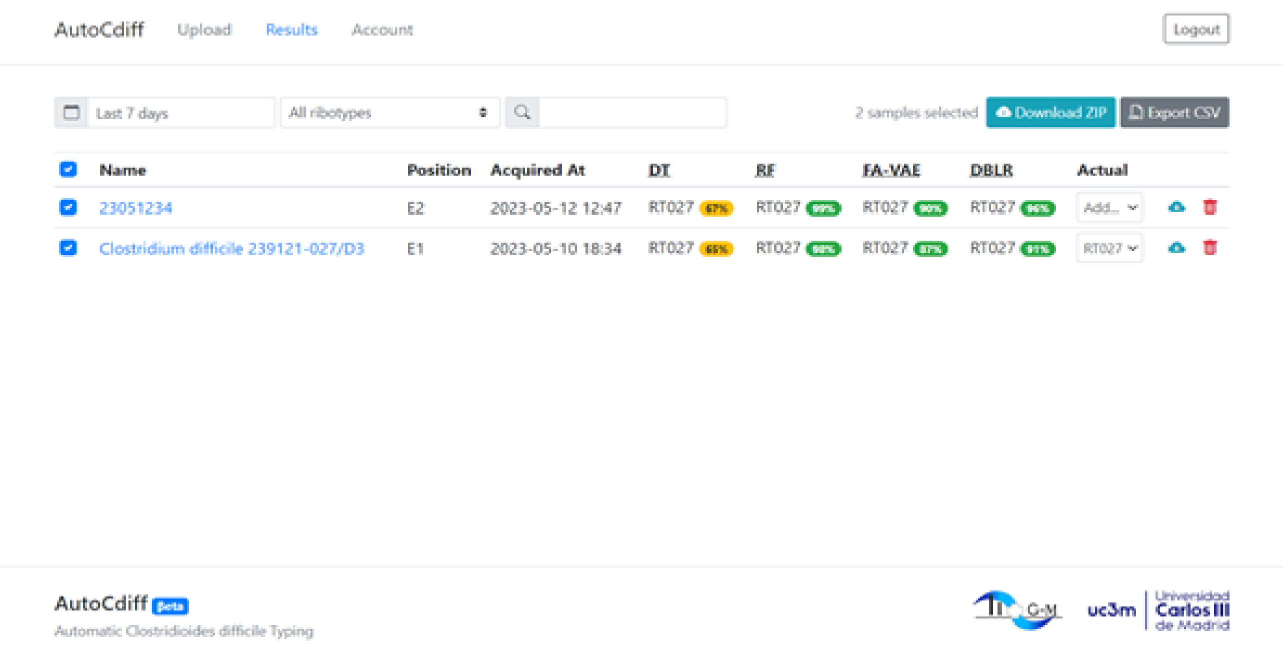


**Figure S4.** Interface of the https://bacteria.id webpage displaying the results provided for 2 MALDI-TOF spectra from suspected RT027 *C. difficile* isolates. The most probable RT is obtained by implementing Decision Trees (DT) and the algorithms RF, FA-VAE and DBLR-FS.
